# Scop3P in 2026: an expanded proteomics-informed resource contextualizing phosphorylation sites through sequence, structure, mutation, and experimental provenance

**DOI:** 10.64898/2026.07.03.736340

**Authors:** Pathmanaban Ramasamy, Natalia Tichshenko, Adrián Díaz, Kevin Velghe, Enrico Massignani, Wim F. Vranken, Lennart Martens

## Abstract

Protein phosphorylation is a central regulatory mechanism controlling protein activity, interactions, and cellular signalling, and its dysregulation is implicated in numerous diseases. Advances in mass spectrometry–based phosphoproteomics have led to a rapid expansion in the number of reported phosphorylation sites; however, interpretation of these data remains challenging due to fragmented evidence, limited structural context, and the lack of uniform experimental provenance across resources. Interpretation is further complicated by the fact that the biological meaning of reported phosphosites can vary substantially across tissues, cell lines, perturbations, and disease settings. Here, we present a major update of Scop3P, a proteomics-informed knowledgebase that contextualizes human phosphorylation sites within integrated sequence, structural, biophysical, evolutionary, and mutational frameworks. The current release incorporates uniformly reprocessed human phosphoproteomics data from 116 PRIDE datasets alongside curated UniProt annotations, retaining peptide-spectrum matches, site localization confidence, and direct links to primary mass spectrometry evidence via Universal Spectrum Identifiers. This integration yields 152,350 unique serine, threonine, and tyrosine phosphorylation sites across 16,533 human proteins, supported by full experimental provenance. Beyond site identification, Scop3P provides residue-level contextual annotations derived from experimentally determined protein structures and proteome-wide AlphaFold models, enabling near-complete structural coverage of phosphorylation sites. Structural context is further complemented by residue-level biophysical, evolutionary, and mutational annotations, supporting integrated assessment of phosphorylation in functional and disease-related settings. The current release also introduces residue interaction network representations derived from AlphaFold-predicted structures, capturing spatial connectivity and local interaction environments of phosphorylation and mutation sites. A redesigned web interface enables interactive exploration through coordinated 1D, 2D, 2.5D, and 3D visualizations, peptide-level coverage views, and direct access to original spectra via PRIDE. By bridging experimental phosphoproteomics with structural, functional, and disease-related context, Scop3P provides a scalable and provenance-aware resource for phosphosite interpretation, hypothesis generation, and data-driven modelling of phosphorylation-dependent regulation.

## Introduction

Protein phosphorylation is one of the most extensively studied post-translational modifications and plays a central role in cellular signalling, protein–protein interactions, and the regulation of protein activity and stability^1–3^. Advances in mass spectrometry-based phosphoproteomics have led to an unprecedented expansion in the number of reported phosphorylation sites (phosphosites) across the human proteome^4–6^. However, the biological interpretation of these sites remains a major challenge. A critical gap persists between the large-scale identification of phosphorylation events and their functional, structural, and mechanistic interpretation^7–9^.

Several widely used resources, including PhosphoSitePlus^10^, dbPTM^11^, and UniProtKB/Swiss-Prot^12^, provide valuable collections of phosphorylation annotations. However, these resources primarily aggregate evidence from heterogeneous sources and often lack direct experimental provenance at the spectrum or experiment level. In addition, the underlying evidence and level of experimental support can vary substantially across annotations, making it difficult to assess the reliability of individual phosphosites^7, 8^. Moreover, the growing number of reported phosphosites across resources does not imply that all annotations are equally well supported, confidently localized, or biologically meaningful^8, 9, 13^. This limitation is further illustrated by our comparison of existing phosphosite resources against the current canonical UniProt reference proteome, which showed that even after restricting datasets to canonical human proteins, a measurable fraction of sites and proteins could not be reconciled with current reference sequences, with the extent of inconsistency varying considerably across resources. Furthermore, access to some widely used phosphosite collections is subject to licensing restrictions that limit redistribution and reuse, whereas others rely primarily on manual extraction of phosphorylation events reported in the literature rather than on systematic reprocessing of the underlying mass spectrometry data, thereby limiting both scalability and the ability to provide direct spectral evidence for individual site identifications.

The biological interpretation of a phosphorylation site is also often context-dependent, varying with the tissue or cell line analysed, the experimental perturbation, the disease setting, and the biological question addressed in the original study^6, 7^. This challenge is particularly relevant for resources that aggregate annotations from heterogeneous literature sources, where phosphosites may be reported without preserving the original experimental and biological context. As a result, experimental phosphoproteomics evidence, structural data, and biophysical context remain insufficiently integrated within a unified, provenance-aware framework^13^. The lack of consistently processed phosphoproteomics evidence and comparable site localization confidence across studies has further hindered comparative analyses and limited the use of these data for downstream modelling^6, 13^.

At the same time, protein structure has emerged as a key determinant of phosphorylation function. The regulatory impact of a phosphorylation event depends not only on the modified residue but also on its spatial location, solvent accessibility, local dynamics, and the conformational state of the protein^14–17^. Historically, the use of structural context has been limited by the scarcity of experimentally determined structures capturing phosphorylated states. Recent advances in deep learning–based structure prediction, most notably AlphaFold^18^, have dramatically increased structural coverage across the proteome^19^. While predicted structures do not capture phosphorylation-induced conformational changes, they nevertheless provide invaluable residue-level spatial context that was previously unavailable at scale.

Scop3P^13^ was originally developed to address these challenges by systematically reprocessing public phosphoproteomics data and integrating phosphosites with protein sequence, structure, and biophysical context. Building on this foundation, the current release substantially expands the resource by combining uniformly reprocessed PRIDE^20^ phosphoproteomics datasets with curated UniProt annotations^12^, explicit localization confidence, and direct links to experimental evidence via Universal Spectrum Identifiers^21^(Figure 1). It further bridges experimental and predicted structural information by integrating both experimentally determined PDB structures^22^ and AlphaFold models^18, 19^, enabling proteome-wide structural contextualization of phosphosites while remaining transparent about data origin and limitations.

**Figure 1.**
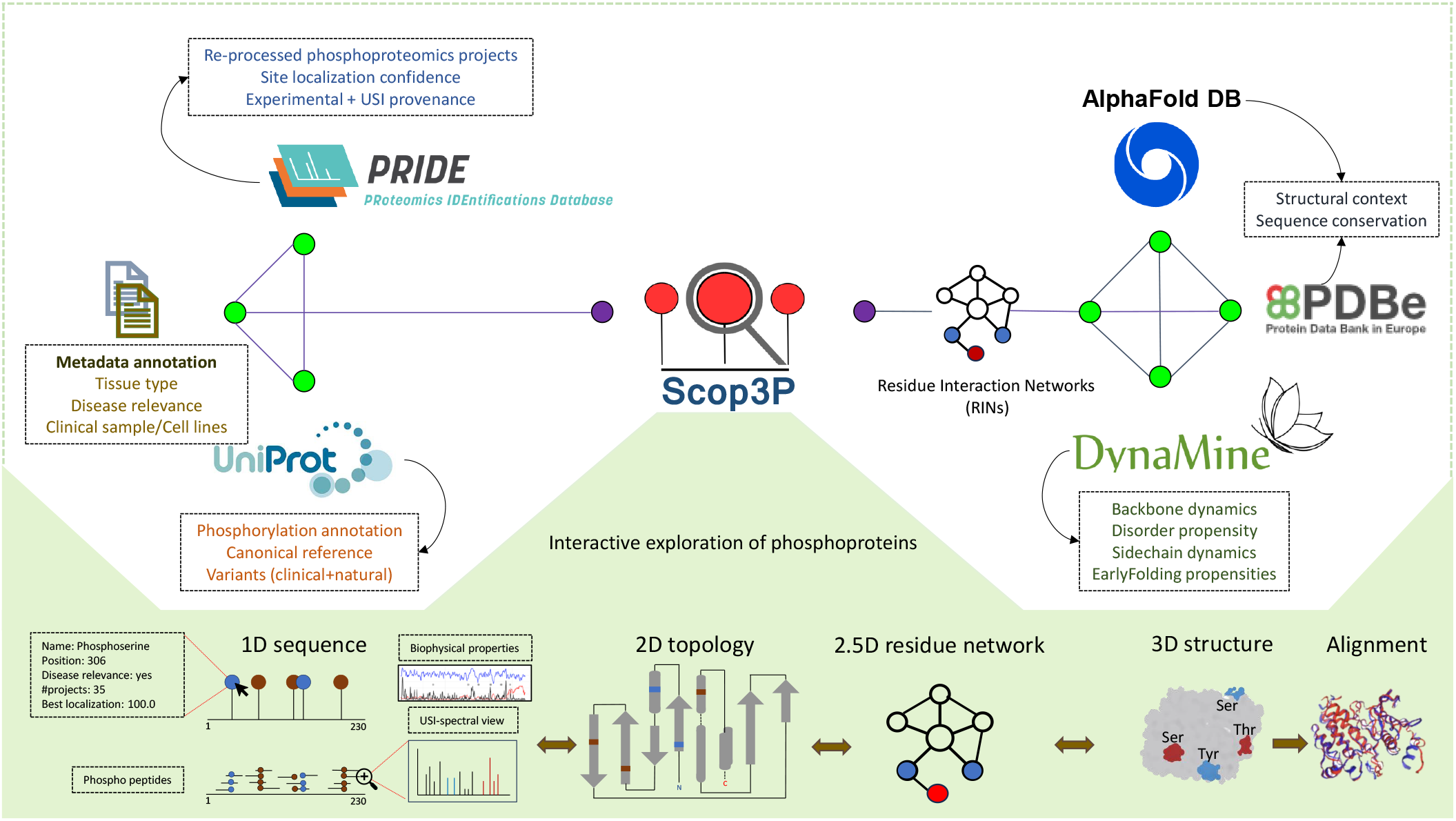
Overview of data sources, annotation layers, and visualization modes in Scop3P. The upper panel shows the integrated data sources: reprocessed phosphoproteomics data with USI provenance from PRIDE, curated annotations and variants from UniProt, predicted structures from the AlphaFold Protein Structure Database, conservation data from PDBe, biophysical predictions from b2btools, and residue interaction networks derived from AlphaFold models. Dataset-level metadata captures tissue type, disease relevance, and sample source. The lower panel illustrates the coordinated 1D, 2D, 2.5D, and 3D visualization modes enabling interactive exploration from sequence to structure.

Beyond site annotation, Scop3P provides residue-level biophysical features, evolutionary conservation, and largescale mutation data mapped onto both experimental and predicted structures. By explicitly capturing gain- and loss-of-phosphorylation-site mutations and distinguishing experimentally observed sites from inferred annotations, Scop3P provides a structure-aware framework for studying phosphorylation, regulatory rewiring, and their perturbation by genetic variation.

### Scop3P 2026 release highlights

Since its original release, Scop3P has expanded substantially in experimental evidence, structural coverage, and contextual annotations. The 2026 release integrates a larger body of uniformly reprocessed phosphoproteomics data, incorporates proteome-wide AlphaFold-predicted structures, expands mutation and disease annotations, and introduces residue interaction network representations, thereby broadening the resource from a phosphosite catalogue to a more context-rich resource for phosphoproteome interpretation. A quantitative comparison between the original release and the 2026 update is summarized in **Table 1**. Beyond expanded phosphosite coverage, the current release adds standardized metadata describing tissue origin, sample source, and disease context across reprocessed datasets, enabling more context-aware interpretation of phosphorylation evidence. These annotations span diverse biological settings, including major cancer types as well as infectious, neurodegenerative, metabolic, and rare-disease contexts.

**Table 1.** Comparison of data statistics between Scop3P 2026 and the previous release.

| Category | Metric | Scop3P 2020 | Scop3P 2026 |
| --- | --- | --- | --- |
| Reprocessed proteomics data | Reprocessed PRIDE datasets | 36 | 116 |
|  | MS/MS runs | 2,016 | 6,823 |
| Phosphoproteome (reprocessed) | Phosphoproteins | 13,510 | 16,297 |
|  | Phosphosites (total) | 93,147 | 140,841 |
|  | Phosphoserine (pS) | 64,009 | 89,856 |
|  | Phosphothreonine (pT) | 21,515 | 34,462 |
|  | Phosphotyrosine (pY) | 7,623 | 16,523 |
| Phosphoproteome (after UniProt integration) | Phosphoproteins | 14,157 | 16,533 |
|  | Phosphosites (total) | 107,894 | 152,350 |
|  | Phosphoserine (pS) | 74,894 | 98,188 |
|  | Phosphothreonine (pT) | 24,025 | 36,568 |
|  | Phosphotyrosine (pY) | 8,975 | 17,594 |
| Evolutionary conservation | Proteins with conservation profiles | 4,384 | 7,717 |
|  | Phosphosites with conservation scores | 14,906 | 87,316 |
| Mutation and disease annotation | Total mutational events | 83,996 | 2,163,169 |
|  | Mutational sites unique | 76,356 | 1,410,744 |
|  | Pathogenic variants | 32,930 | 203,934 |
|  | Mutational events occurring at phosphosites (pSTY → any) | 680 | 29,172 |
|  | Loss of STY sites (STY → non-STY) | 11,000 | 283,901 |
|  | Gain of STY sites (non-STY → STY) | 12,979 | 314,654 |
|  | Preserved STY sites (STY → STY) | 1,209 | 39,724 |
| Structural coverage | Proteins with experimental PDB structures | 4,384 | 7,685 |
|  | Phosphosites mapped to experimental PDB structures | 14,906 | 25,284 |
|  | Proteins with predicted AlphaFold models | 13,854 | 19,266 |
|  | Phosphosites mapped to AlphaFold structures | 103,100 | 147,340 |
|  | Mutation sites mapped to experimental structures | 27,247 | 379,787 |
|  | Mutation sites mapped to AlphaFold structures | 53,385 | 1,310,563 |

In parallel, the web interface has been redesigned to support multiscale exploration of phosphorylation data. At the sequence level, the original lollipop-style phosphosite representation is retained in the protein summary view, where it conveys site-level reliability based on identification confidence across datasets. This is complemented by a track-based visualization using the Nightingale web components^23^, enabling the simultaneous display of phos-phorylation sites together with functional domains, biophysical properties, and sequence variants within a unified sequence context. This integrated view allows users to interpret individual sites in relation to neighboring residues and overlapping annotations. Protein-level peptide coverage visualizations further extend this sequence-centered representation by providing an overview of phosphorylated peptides identified for a given protein across reprocessed datasets, thereby helping users assess the experimental coverage of phosphorylated regions.

Beyond the sequence level, two-dimensional topology diagrams provide a simplified overview of secondary structure organization, using a topology viewer adapted from the framework developed by PDBe^24, 25^ and here extended to support PTM-specific annotations and interactive exploration. Residue interaction network representations provide an additional layer of structural context by capturing residue-level contacts and spatial proximity, allowing users to examine how phosphorylation and mutation sites are embedded within their local structural environment. Three-dimensional molecular visualization is now provided using Mol*^26^, replacing the NGL viewer^27^ used in the original release. The updated viewer also supports structure alignment, enabling users to superimpose pairs of structures, including experimental and AlphaFold-predicted models or modified and unmodified conformations, to assess conformational differences in the context of annotated phosphorylation sites. Together, these coordinated 1D, 2D, 2.5D, and 3D views link experimental provenance with sequence, structural, and network-level context (Figure 1, Figure 3, Figure 4, Figure 5).

**Figure 2.**
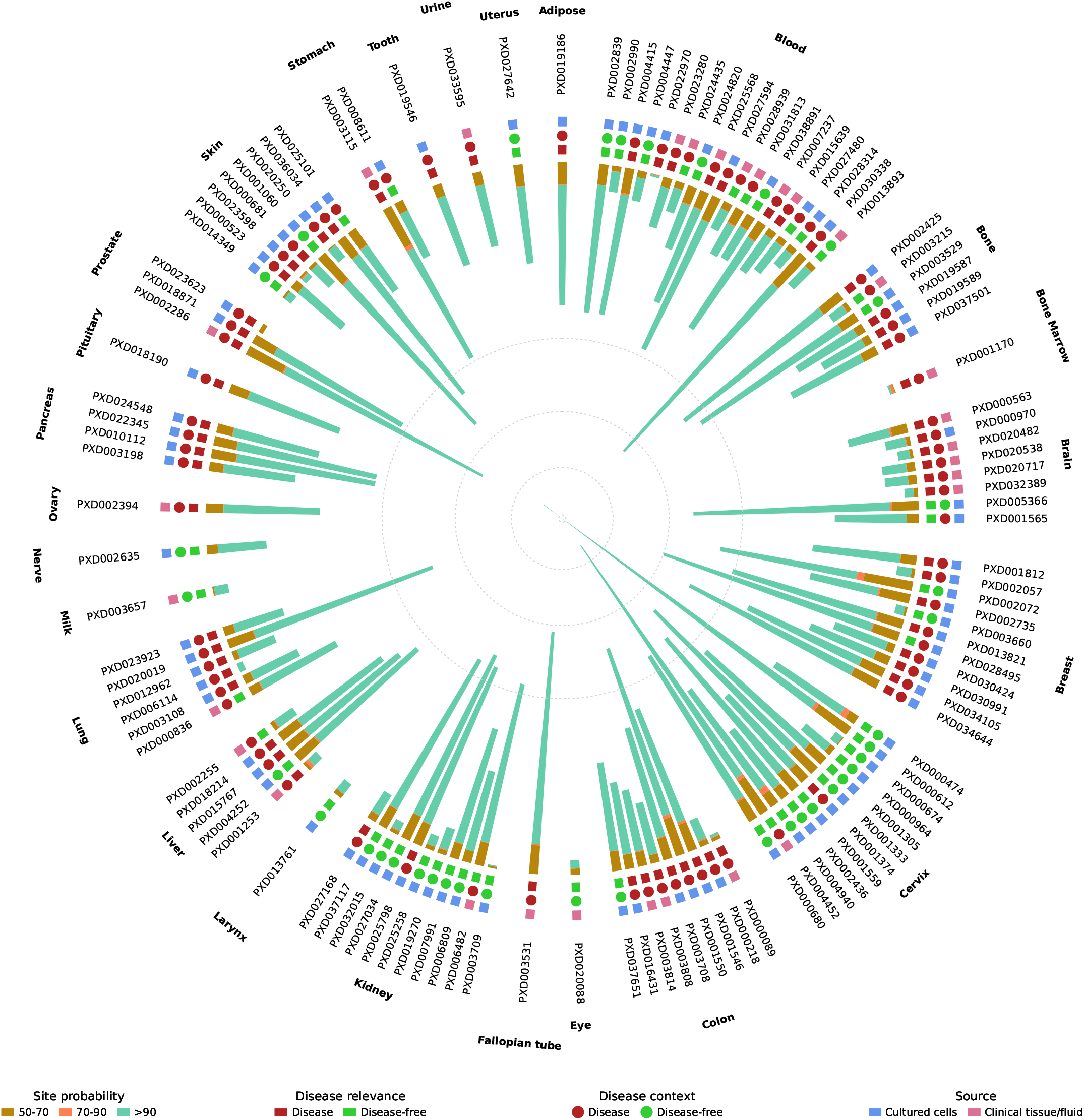
Expansion and contextualization of reprocessed phosphoproteomics data in Scop3P. Circular overview of PRIDE datasets grouped by tissue or sample origin. Radial bars represent dataset-specific phosphosite counts and their localization probability, while colored tiles encode disease relevance, inferred disease context, and sample source. The figure illustrates the increased dataset coverage and integration of biological and disease metadata across diverse experimental settings.

**Figure 3.**
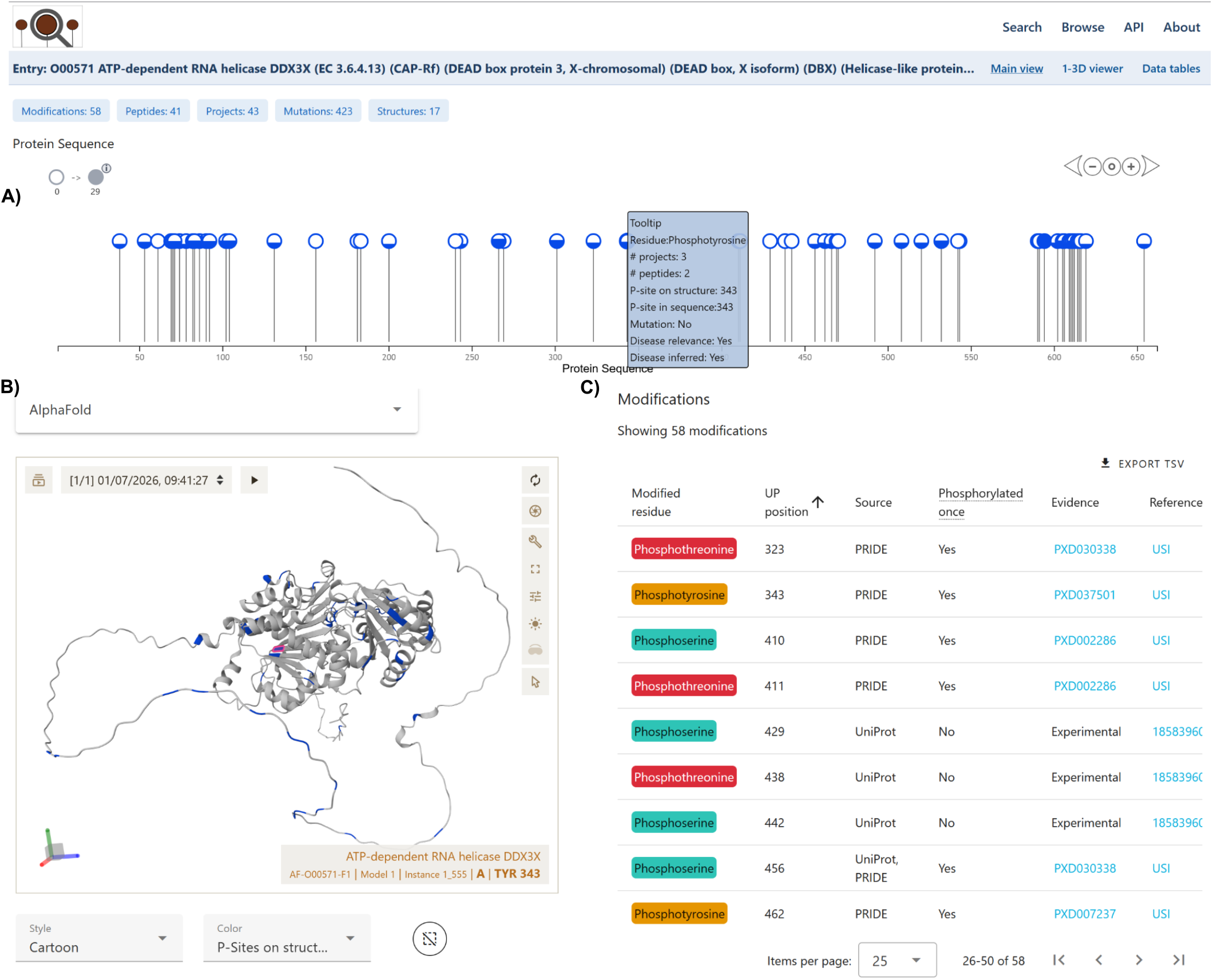
Sequence and structure-level exploration of a phosphoprotein in the Scop3P web interface, illustrated for ATP-dependent RNA helicase DDX3X (UniProt O00571). (A) Protein-level lollipop plot of all identified modifications; ball size and fill encode identification frequency across projects, and hovering over a site shows its metadata (residue type, supporting peptides and projects, structural mapping, mutation status at the site, disease context inferred and disease relevance derived from LLM-assisted, manually validated metadata curation) while highlighting the corresponding residue on the structure in (B). (B) AlphaFold-predicted structure in Mol*, with the selected modification highlighted and selectable models, representation styles, and colouring schemes. (C) Modifications table listing modified residue, UniProt position, evidence source, and reference; USI links connect directly to the PRIDE spectrum viewer, linking each site to its original mass spectrometry evidence.

**Figure 4.**
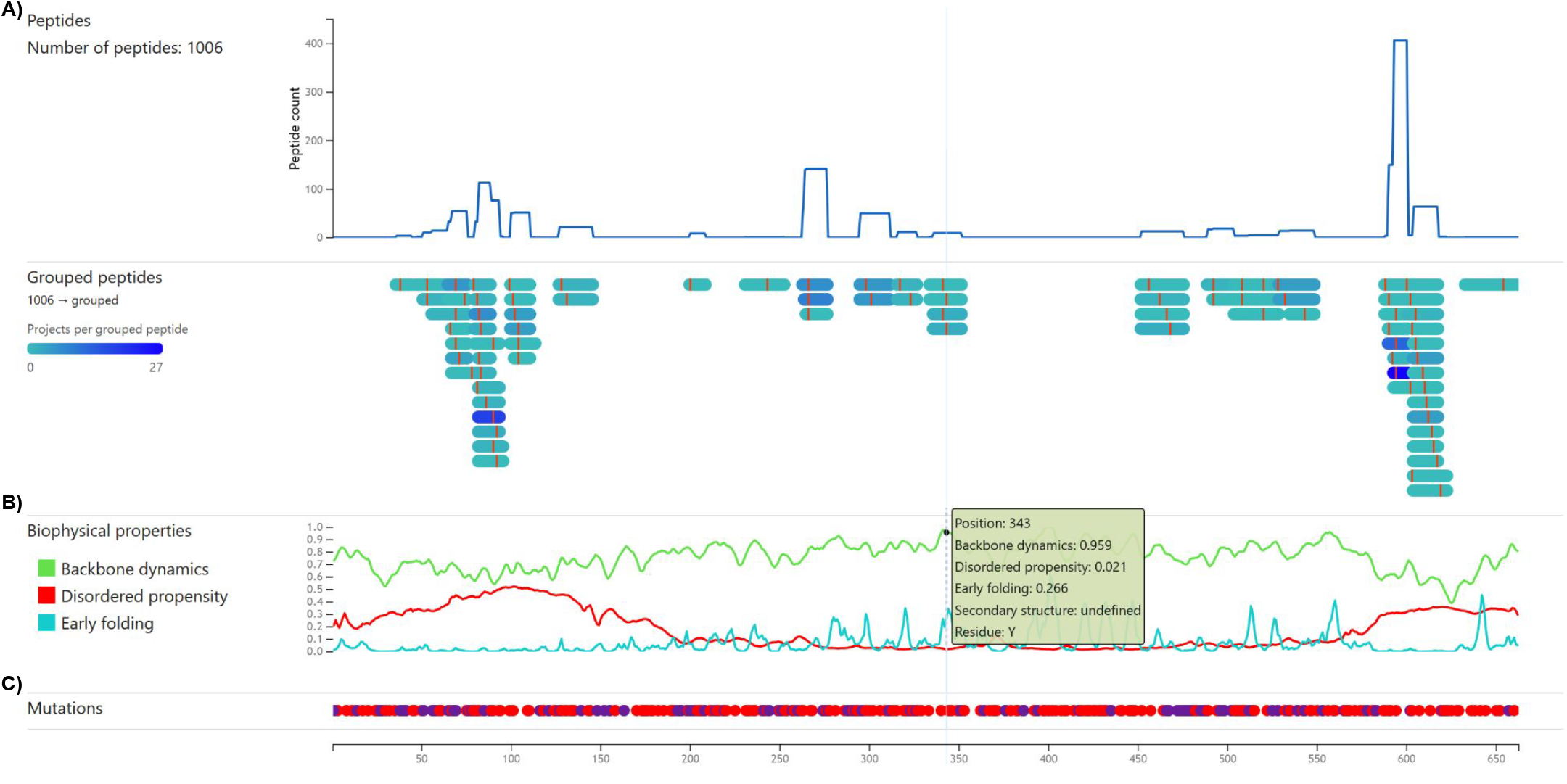
Residue-level annotation tracks along the protein sequence in the Scop3P web interface, shown for the same phosphoprotein as in Figure 3 (DDX3X, UniProt O00571). (A) Phosphopeptide coverage plot: number of identified phos-phopeptides per sequence position (top) and grouped peptide tracks coloured by number of contributing projects (bottom), indicating regions of strongest experimental support. (B) Residue-level biophysical property tracks (backbone dynamics, disorder propensity, and early-folding propensity, predicted by DynaMine, DisoMine, and EfoldMine). (C) UniProt-derived mutation track along the sequence.

**Figure 5.**
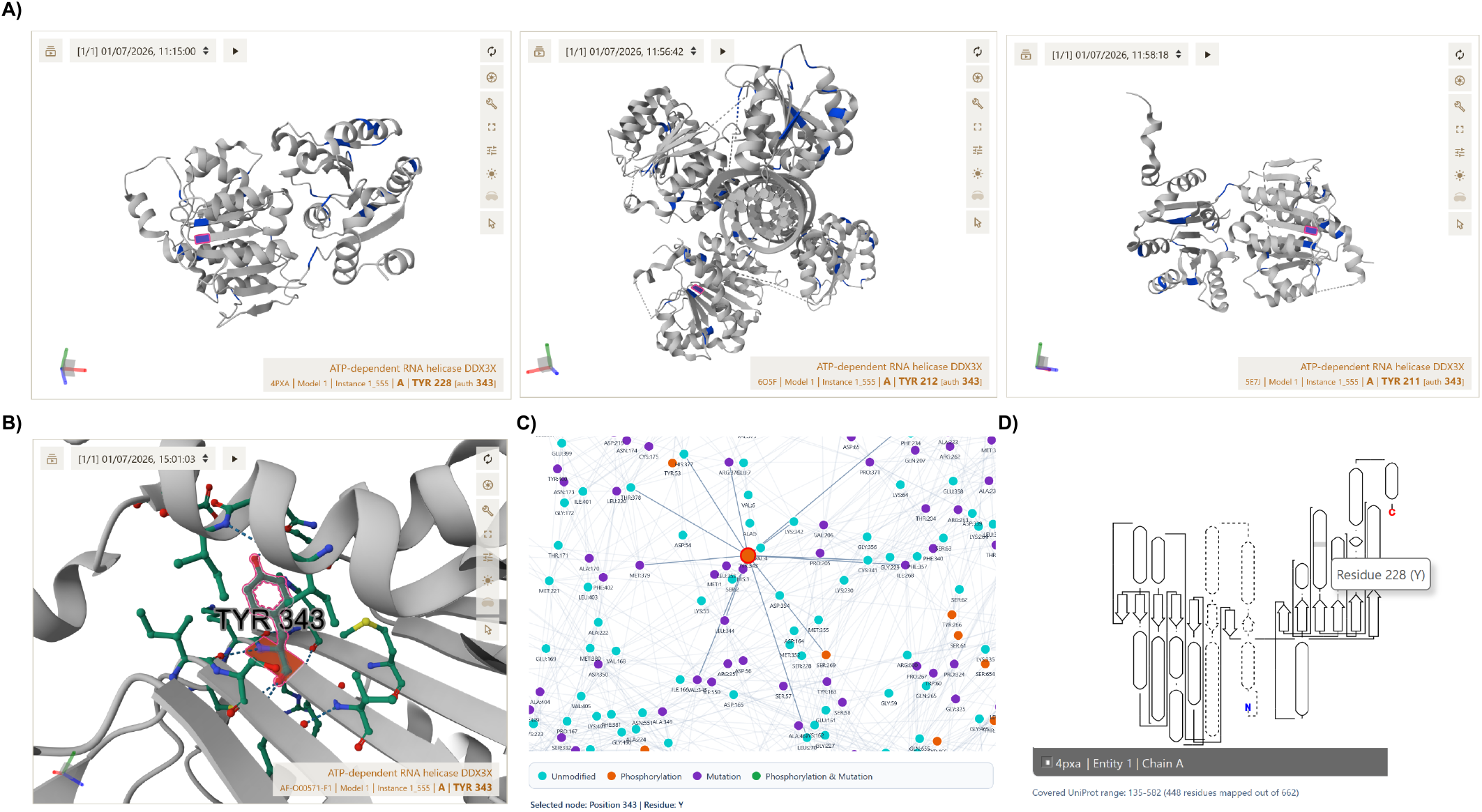
Structural and network-level context of a single phosphosite in Scop3P, shown for TYR343 of DDX3X. (A) The same site mapped onto three independent experimentally determined PDB structures (4PXA, 6O5F, 5E7J; author-numbered 228, 212, and 211, respectively), illustrating structural context across multiple deposited entries. (B) AlphaFold model zoomed to the modified residue and its neighbouring side chains, showing local contacts. (C) Residue interaction network centred on the site, with connected residues coloured by status (unmodified, phosphorylation, mutation, or both). (D) 2D topology diagram of the PDB structure shown in (A), with the corresponding residue highlighted.

### Expansion of uniformly reprocessed phosphoproteomics data

Scop3P integrates uniformly reprocessed large-scale human phosphoproteomics datasets deposited in PRIDE^20^, substantially expanding the experimental foundation of the resource. The current release includes 116 reprocessed PRIDE datasets comprising 6823 raw mass spectrometry runs, in which approximately 192.56 million MS/MS spectra were searched (Figure 2).

All datasets were reanalysed using the target-decoy workflow established in the original release of Scop3P^13^, with results filtered at 1% false discovery rate (FDR). To account for differences in experimental design, raw files were assigned to search groups defined by enzyme, labelling strategy, and variable modifications, and searched using harmonized parameters within each group (Supplementary Table S1). Across all groups, carbamidomethylation of cysteine was set as a fixed modification, while phosphorylation of serine, threonine, and tyrosine, N-terminal acetylation, and oxidation of methionine were included as common variable modifications.

At a stringent PSM-level threshold of q ≤ 0.01, this yielded approximately 7.36 million phospho-PSMs. Phosphorylation site localization probabilities were computed using PhosphoRS^28^, enabling site-level confidence assessment across all identifications. Applying a localization filter of at least 0.50 retained approximately 5.73 million phospho-PSMs, corresponding to 242,800 phosphopeptidoforms and 125,683 unique peptide backbones. These mapped to 140,841 unique serine, threonine, and tyrosine phosphosites across 16,297 proteins, including 117,832 sites with localization probability greater than 0.90 (Figure 2).

Phosphosites were retained when supported by at least one identification with a localization probability of 0.50 or higher. All supporting experiment, peptide, and PSM-level evidence remained accessible in the detailed data tables. When combined with curated UniProt^12^ phosphorylation annotations, this experimental core expands to 152,350 unique serine, threonine, and tyrosine phosphosites across 16,533 human proteins, reflecting the complementary contributions of large-scale experimental reprocessing and expert curation (**Table 1**). The UniProt-derived phos-phorylation and variation annotation layers that contribute to this integrated resource are described in detail in the section “Mutation and phosphorylation context from UniProt”.

To quantify annotation consistency across major public phosphosite resources, we mapped entries from Phospho-SitePlus^10^, dbPTM^11^, and the reprocessed phosphoproteomics compendium of Ochoa et al.^6^ against the current canonical UniProt reference proteome (Supplementary Table S2). After restricting all datasets to canonical human proteins, dbPTM^11^ showed the highest inconsistency, with 4.6% of sites and 10.3% of proteins failing validation against the current canonical reference, compared with 1.1% of sites and 1.4% of proteins for PhosphoSitePlus^10^, and 2.1% of sites and 2.1% of proteins for the Ochoa dataset^6^. Together, these results highlight the extent to which phosphosite resources can differ in reference consistency even when they draw on proteomics-derived evidence. Although several of these resources incorporate proteomics annotations, their evidence is generally aggregated from independently processed datasets without harmonized search parameters, comparable localization scoring, or direct links to the underlying mass spectra. Scop3P addresses these limitations by uniformly reprocessing raw mass spectrometry data, anchoring all sites to the current canonical UniProt reference proteome^12^, and exposing spectrum-level provenance through Universal Spectrum Identifiers^21^. All data are distributed under a CC BY 4.0 license, ensuring unrestricted reuse.

For summary-level representations, including protein-centric views and one-dimensional site plots, a single representative peptide, localization score, and USI were selected for each site using the highest localization probability observed across all supporting identifications and datasets. These summaries display the best-supported observation for each site while preserving direct links to the originating PRIDE datasets and, where available, to the underlying mass spectra, ensuring transparent and reusable experimental provenance (Figure 3C).

To support interpretation of these phosphorylation data beyond site counts alone, the reprocessed datasets were further annotated with standardized metadata describing tissue origin, sample source, and disease-related context, thereby preserving the biological setting in which phosphosites were originally observed.

### Metadata extraction and disease context annotation

To systematically characterize the biological and clinical context of the collected phosphoproteomics datasets, we developed a standardized metadata curation framework based on the primary literature. For each PRIDE dataset (PXD), associated source materials, including full-text articles, supplementary information, and PRIDE^20^ API annotations where available, were examined to extract information on sample origin, experimental design, and biological interpretation. Initial metadata extraction was assisted by a large language model (LLM), which was used to organize key study attributes such as tissue type, specific tissue of origin, cell type or cell line identity, enrichment strategy, and disease-related study aims. All LLM-derived annotations were subsequently reviewed and manually verified against the original source materials using predefined curation criteria to ensure accuracy and consistency. This metadata layer was designed to preserve the experimental and biological context in which phosphosites were originally reported, including tissue or cell model, study design, and disease relevance.

Across the 116 reprocessed datasets, metadata curation captured substantial biological diversity in sample origin and study context (Supplementary Table S3). The collection included 86 datasets based on cultured cells and 30 based on clinical tissue or fluid samples, spanning a broad range of tissues, with blood, breast, cervix, colon, and kidney among the most frequently represented groups. The annotated studies also covered diverse disease settings, including multiple cancer types such as colorectal, cervical, breast, prostate, pancreatic, and hematological malignancies, as well as neurodegenerative, infectious, metabolic, and rare-disease contexts, including Alzheimer’s disease, COVID-19, HCMV infection, malaria-related models, non-alcoholic steatohepatitis, type 2 diabetes, and rare syndromic disorders. Disease-context annotation identified 65 datasets in which both the phosphosite-level interpretation and the broader experimental context were classified as disease-related, 37 datasets classified as disease-free in both respects, and 14 datasets in which phosphosites were not considered directly disease-relevant despite being derived from models with inferred disease context. This distribution highlights the importance of separating sample origin and model background from phosphosite-level biological interpretation.

The curation framework was designed to distinguish sample origin from the disease relevance of the phospho-proteomic observations. For each dataset, we annotated the broad sample class, such as cultured cells, primary cells, clinical samples, or animal models, together with the specific tissue, cell line, or experimental system used. Disease conditions were extracted from the source materials and standardized using Disease Ontology identifiers (DOID) where applicable. Additional experimental metadata, including phosphopeptide enrichment strategy, mass spectrometry instrumentation, and relevant study notes, were also recorded to support downstream evaluation of dataset quality and reusability (Supplementary Table S3).

A key feature of the annotation strategy was the distinction between phosphosites that were meaningfully linked to disease biology and those merely detected in disease-derived material. To capture this, two complementary metadata fields were introduced: *Disease_relevance_of_phosphosites* and *Disease_context_inferred. Disease_relevance_of_ phosphosites* was assigned as positive only when the experimental design explicitly investigated a disease mechanism, pathogenic perturbation, biomarker question, or therapeutic response, and when the identified phosphosites were interpreted in that context. Accordingly, datasets focused on methodological benchmarking, analytical optimization, generic signaling studies, or basic cell biology were not classified as disease-relevant at the phosphosite level, even when disease-derived cell lines were used, preventing disease associations from being inferred from sample origin alone.

The *Disease_context_inferred* field captured datasets that could still inform disease biology despite not being explicitly framed as disease-focused phosphoproteomics studies. This included datasets based on patient tissues, patient-derived models, pathogenic infection systems, genetically or chemically perturbed models designed to mimic clinical pathology, and studies comparing distinct disease states. By contrast, the routine use of standard cancer cell lines as generic experimental systems did not automatically qualify as inferred disease context unless the model itself was central to a clinically relevant biological question.

For datasets lacking a corresponding full-text publication, annotations were derived from PRIDE API-derived dataset descriptions, sample metadata, and data processing protocol annotations. Where the available information was insufficient to determine sample origin, enrichment strategy, or biological context with confidence, the dataset was classified as insufficient data (Supplementary Table S3). Together, this LLM-assisted and manually validated workflow generated a high-confidence metadata layer that distinguishes phosphosites directly linked to disease pathogenesis or treatment response from those identified in baseline, methodological, or broadly mechanistic contexts.

### Structural coverage and conformational context

#### Experimentally determined structures

Scop3P integrates phosphosites with experimentally determined protein structures deposited in the Protein Data Bank (PDB)^22^, providing direct three-dimensional context wherever available. In total, 7,685 human proteins in the database are associated with at least one experimentally resolved structure, corresponding to 64,066 unique PDB entries and 149,063 PDB chain combinations.

Despite the breadth of structural data in the PDB, experimental coverage of phosphosites remains limited. Of the 152,350 unique phosphosites represented in the current release, only 25,284 sites, approximately 16%, can be mapped to at least one experimentally determined structure. This highlights a central challenge in the field: although phosphorylation is widespread, structures capturing the relevant regions, and especially the modified states, remain comparatively rare.

Among phosphosites with structural coverage, only a small subset is captured in an experimentally modified state. Specifically, 484 unique sites are resolved as phosphorylated residues in PDB structures, including 232 SEP, 119 TPO, and 133 PTR, underscoring the limited availability of experimentally determined phospho-specific structural information. All valid site-structure associations were retained, enabling comparison across modified and unmodified structural contexts where available. Structural coverage is also often not restricted to a single entry, with the median phosphosite represented in three distinct PDB structures, allowing users to examine many sites across multiple experimentally observed structural contexts.

Structural quality is further reflected in the resolution metadata of linked PDB entries: 79% of associated structures are resolved at 3 Å or better, providing a large set of high-confidence experimental models for downstream structural analyses.

#### Integration of AlphaFold-predicted structures

To address limited experimental coverage, the current release systematically integrates predicted protein structures from the AlphaFold Protein Structure Database^29^. AlphaFold models are available for 19,266 human proteins and provide near-complete structural coverage across the proteome. When AlphaFold structures are included, structural context can be assigned to approximately 97% of all phosphosites, dramatically expanding coverage relative to experimental structures alone.

AlphaFold-derived models were used to compute secondary structure and solvent accessibility using DSSP^30^, ensuring methodological consistency with PDB-based annotations. Predicted structures are explicitly distinguished from experimentally determined models, as AlphaFold predictions typically represent a dominant conformational state and do not capture phosphorylation-induced structural rearrangements^31^. By integrating PDB and AlphaFold structures, Scop3P provides a practical balance between experimental realism and proteome-wide coverage, while remaining transparent about the origin and limitations of each annotation. In addition to expanding residue-level structural coverage, AlphaFold models provide the structural basis for deriving residue interaction network representations across nearly the entire human phosphoproteome.

### Residue interaction network context

To further characterize the spatial environment of phosphorylation and mutation sites, Scop3P incorporates residue interaction network (RIN) representations derived exclusively from AlphaFold-predicted models^18, 29^. In these networks, residues are represented as nodes connected by spatial proximity, providing an explicit framework for capturing local interaction environments and residue-level structural connectivity.

Residue interaction networks were constructed on a per-chain basis, with edges defined between residue pairs whose C*α* or C*β* atoms lie within 8 Å^32^. Networks were generated separately using either C*α*–C*α* or C*β*–C*β* distance definitions; for glycine residues, C*α* atoms were used in place of C*β* atoms. Edge weights were assigned using an inverse-distance scheme, such that closer residue pairs contribute more strongly to network connectivity^33, 34^. To account for structural uncertainty in AlphaFold models, edge weights were further scaled by the per-residue pLDDT confidence scores, reducing the contribution of interactions involving low-confidence regions.

RIN-based annotations enable identification of phosphorylation and mutation sites located in densely connected or structurally central regions, complementing linear sequence features such as evolutionary conservation, solvent accessibility, and secondary structure (Figure 5C). This network-level perspective captures spatial relationships that are difficult to infer from sequence or static structural representations alone. In particular, it enables exploration of the spatial proximity between phosphorylation and mutation sites, including direct overlap and connectivity within local structural neighborhoods.

### Mutation and phosphorylation context from UniProt

In addition to uniformly reprocessed phosphoproteomics evidence, Scop3P integrates curated phosphorylation and mutation annotations retrieved directly from the UniProt^12^ Proteins API. This UniProt-derived annotation layer complements the experimental reprocessing workflow, contributes to the final integrated counts of phosphosites and phosphoproteins reported above, and situates experimentally observed sites within a broader functional and disease-related framework. Phosphorylation annotations were extracted from the UniProt^12^ feature set, retaining experimentally supported and curator-reviewed serine, threonine, and tyrosine modification events recorded using canonical UniProt residue numbering. Associated evidence codes and literature references were preserved to maintain annotation provenance.

In parallel, sequence variants with reported clinical significance were retrieved from UniProt^12^ variation annotations. Each variant was recorded with its affected residue position, wild-type and mutated amino acid, consequence type, clinical significance category, and supporting evidence. Phosphosites and clinically annotated variants were then mapped onto both experimentally determined and AlphaFold-predicted structures^18, 29^ using a unified residue-mapping pipeline, ensuring consistent integration of post-translational modification and mutation data.

Here, mutational events denote distinct amino acid substitutions recorded at a residue position, whereas mutation sites denote unique residue positions that may harbor one or more such events. In total, Scop3P includes nearly 2.16 million mutational events across 18,246 human proteins. When considering experimentally determined structures alone, approximately 27% of mutation sites can be mapped to at least one PDB structure^22^. This coverage increases substantially when AlphaFold models^18, 29^ are included, with over 92% of mutation sites associated with predicted structures. This integrated view allows phosphorylation and clinically annotated variation to be examined within a shared sequence and structural framework.

A particularly informative subset of variants directly intersects with phosphosites. Scop3P identifies 29,172 unique mutation events that coincide with experimentally observed phosphosites, representing positions at which genetic variation may directly disrupt phosphorylation-dependent regulation. In addition, mutations that introduce serine, threonine, or tyrosine residues are captured as potential gain-of-phosphorylation-site events, providing insight into regulatory rewiring (Table 1). Together, these annotations enable residue-level interpretation of the interplay between phosphorylation and genetic variation, even when experimentally resolved phospho-specific structures are unavailable.

### Residue-level evolutionary conservation

The current release of Scop3P integrates residue-level conservation profiles obtained from the PDBe Graph API^24, 35^, replacing the AMINODE^36^ derived conservation scores used in the previous version. The PDBe API provides position-specific amino acid probability distributions derived from multiple sequence alignments of structurally characterized homologues, offering finer-grained conservation information than the single summary scores previously available. For each protein, the canonical UniProt^12^ sequence was used as the reference coordinate system. Conservation data were mapped to UniProt residue positions using the index-to-sequence correspondence provided in the PDBe^24^ payload, and the probability corresponding to the observed amino acid at each position was assigned. Positions lacking a corresponding mapping in the PDBe alignment^24^ were recorded as missing. Probability values were scaled to a 0–100 range and stored as residue-level annotations aligned to UniProt coordinates. This conservation layer enables direct comparison with phosphorylation, mutation, and structural features within a unified sequence framework, facilitating the identification of conserved regulatory sites and regions of potential functional importance.

### System architecture, visualization framework, and API endpoints

The current release of Scop3P was implemented using Vue.js 3 (https://vuejs.org) as the frontend framework, with Vuetify as the component and styling library. Interactive visualizations were primarily developed in D3.js^37^ to allow flexible rendering and customized user interaction. The backend was rebuilt in Python using FastAPI (https://fastapi.tiangolo.com) for API management, SQLAlchemy (https://www.sqlalchemy.org) for database interaction, and Pydantic for schema definition and data validation. Data are stored in a PostgreSQL 16 relational database (https://www.postgresql.org).

The visualization layer integrates several complementary technologies to support sequence and structure-level exploration of phosphosites. Three-dimensional protein structures are rendered using Mol*^26^ (Figure 3B, Figure 5A,B). Two-dimensional topology diagrams were adapted from the PDBe^24, 35^ topology viewer and extended to support Scop3P-specific PTM annotation and user interaction. Sequence-based visualization retains the original lollipop-style phosphosite representation along the primary amino acid sequence, together with the concentric-ring biophysical plots from the first release. In these plots, amino acids occupy the inner ring, while residue-level properties, including backbone dynamics and disorder propensity, are displayed in the outer rings. These views are further complemented by integration of the Nightingale web components^23^, which provide a comprehensive track-based representation of phosphosites, domains, mutations, and biophysical features within a unified sequence context (Figure 4B). Residue interaction network visualizations were implemented in D3.js^37^ using force-directed layouts to represent residue-level contacts and spatial relationships (Figure 5C). In addition, the structural visualization module supports pairwise structure alignment for superposition-based comparison of experimental and predicted structures, as well as modified and unmodified conformations.

Programmatic access is provided through REST API endpoints that expose protein, phosphosite, peptide, and experiment-level annotations in machine-readable form. These endpoints support retrieval of phosphorylation evidence, localization confidence, sequence and structure mappings, and associated provenance metadata, thereby facilitating integration with downstream computational workflows and external resources.

### Web interface and data access

Scop3P provides a web-accessible interface for large-scale, residue-centric exploration of phosphorylation evidence and contextual annotations. Three-dimensional structural visualization is provided using Mol*^26^, enabling consistent rendering of both experimentally determined and AlphaFold-predicted structures within the same exploratory framework. The interface supports interactive exploration of phosphosites, mutations, and biophysical annotations through coordinated 1D, 2D, 2.5D, and 3D views, allowing users to move seamlessly from experimental evidence to sequence-level and structural interpretation. Protein-level summaries are linked to detailed peptide- and experiment-level evidence tables, enabling direct inspection of localization confidence, supporting peptides, associated PRIDE datasets, and linked spectra where available. This design allows users to trace individual phosphosites from high-level overview to underlying experimental provenance and structural context.

Scop3P integrates both sequence-level and structure-level information for human phosphoproteins and individual phosphosites. All annotations are mapped onto human protein sequences obtained from UniProtKB/Swiss-Prot^12^, with sequence-to-structure residue mapping performed using SIFTS^38, 39^. Phosphosites located in unresolved or missing regions of available structures are excluded from structure-based analyses. Residues with structural information are annotated with structural and evolutionary features as described above. In addition, residue-level biophysical properties, including backbone dynamics, disorder propensity, and early folding propensity, are predicted using the b2bTools suite^40^, which integrates DynaMine^41^, DisoMine^42^, and EfoldMine^43^, are mapped onto the corresponding protein sequences (Figure 4B).

To estimate phosphosite reliability, each site is annotated based on its frequency of identification across phosphoproteomics experiments. As a second layer of evidence, each phosphosite is linked to the number of distinct peptides identified for the corresponding protein across PRIDE datasets, and each peptide is further annotated with its ProteomeXchange^44^ identifier and its frequency of detection across datasets.

Scop3P is designed to map all phosphosites of a given protein onto all available three-dimensional structures. When multiple structures are available, all structures containing at least one mapped phosphosite are retained for visualization and annotation (Figure 5A). For phosphosites with structural information, additional annotations include assembly and interface context, macromolecular chain assignment, secondary structure, and relative solvent accessibility derived from DSSP-based^30^ analyses.

Scop3P is distributed under the Creative Commons Attribution 4.0 (CC BY 4.0) license, facilitating free reuse with appropriate attribution. Together with explicit experiment-level metadata and Universal Spectrum Identifiers (USIs)^21^, this framework ensures that phosphorylation data in Scop3P adhere to FAIR (Findable, Accessible, Interoperable, and Reusable) principles while preserving clear attribution to the original data producers and associated PRIDE datasets.

## Discussion and future directions

Scop3P 2026 reflects both the progress and the limitations of contemporary phosphoproteomics and structural biology. Although the number of experimentally resolved phospho-dependent conformational states remains small, the depth and quality of experimental evidence supporting phosphorylation site identification continue to improve. By preserving provenance, integrating diverse contextual annotations, and explicitly representing uncertainty and sparsity, Scop3P provides a realistic and reusable foundation for future studies.

A key strength of the current release is that phosphosites are not treated as universally interpretable entities, but as observations whose functional meaning may depend on tissue context, cell model, perturbation, disease setting, and experimental design. This context-aware representation is particularly important for large phosphosite collections assembled from heterogeneous studies, where the same residue may be observed under markedly different biological conditions.

Beyond interactive exploration, Scop3P is well suited for the development and benchmarking of data-driven models aimed at predicting phosphorylation site function, regulatory importance, and conformational effects. The combination of spectral evidence, localization confidence, evolutionary conservation, structural context, and mutation data is difficult to assemble from isolated sources, making Scop3P a valuable benchmark resource for computational modelling.

Future updates will focus on expanding contextual annotations that place phosphosites in their biological setting, including tissue and cell-type specificity, subcellular localization, and disease relevance. In parallel, Scop3P will incorporate additional model organisms supported by high-quality proteomics data in PRIDE and will continue to integrate emerging experimental and predicted structural information as it becomes available. Future releases will also extend residue interaction network analysis to experimentally determined PDB structures, enabling direct comparison between predicted and experimentally resolved interaction networks and facilitating the analysis of phosphorylation-dependent conformational changes where multiple structures are available. Together, these developments will further establish Scop3P as a provenance-aware, context-rich resource for interpreting phosphorylation across structural, biological, and disease-related contexts.

**Supplementary Table S2.**
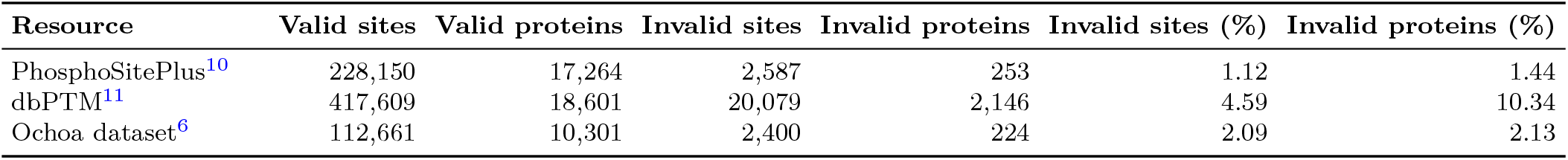
Canonical UniProt sequence mapping statistics for the integrated PTM resources. Valid entries were successfully mapped to the canonical UniProt protein sequence, whereas invalid entries could not be mapped. Percentages represent the fraction of invalid entries relative to the total number of entries (valid + invalid).

| Resource | Valid sites | Valid proteins | Invalid sites | Invalid proteins | Invalid sites (%) | Invalid proteins (%) |
| --- | --- | --- | --- | --- | --- | --- |
| PhosphoSitePlus<sup>10</sup> | 228,150 | 17,264 | 2,587 | 253 | 1.12 | 1.44 |
| dbPTM<sup>11</sup> | 417,609 | 18,601 | 20,079 | 2,146 | 4.59 | 10.34 |
| Ochoa dataset<sup>6</sup> | 112,661 | 10,301 | 2,400 | 224 | 2.09 | 2.13 |

## Supplementary Information

The following supplementary material accompanies this manuscript:

- **Supplementary Table S1**. Search group definitions and harmonized database search parameters used for uniform reprocessing of PRIDE phosphoproteomics datasets.
- **Supplementary Table S2**. Canonical UniProt sequence mapping statistics for the integrated PTM resources.
- **Supplementary Table S3**. Curated metadata annotations for all uniformly reprocessed PRIDE phosphoproteomics datasets, including sample origin, biological context, disease annotation, experimental design, phosphopeptide enrichment, and mass spectrometry metadata.

## Acknoweldgments

The authors thank all data submitters to the PRIDE database for making their proteomics data publicly available, and the developers of the external resources used in this study for providing open-access data and API platforms. P.R., A.D., and N.T. thank the participants of BioHackathon Europe 2025 for their contributions. P.R. and A.D. thank the participants of the Scop3P training sessions for their valuable feedback. P.R. thanks the members of the CompOmics and Bio2Byte groups for feedback on the Scop3P platform.

## Author Contributions

P.R., L.M., and W.F.V. conceived the project. P.R. designed the study, developed the overall system architecture, and performed the data acquisition, reprocessing design, annotation, curation, and integration. N.T. led the front-end development of the web interface, developed the visualization components, and led the development of the Python API client. A.D. contributed to the development of interface components and to the packaging and distribution of the Python API client. E.M. and K.V. carried out the phosphoproteomics reprocessing. P.R., A.D., and N.T. contributed to workflow design and software development. P.R., W.F.V., and L.M. supervised the project and acquired funding. All authors contributed to manuscript preparation and approved the final version.

## Conflict of Interest

The authors declare no competing interests.

## Funding

This work was supported by a BOF grant from Ghent University [BOF/01P07523 to P.R.], by the Research Foundation-Flanders (FWO) [G028821N to P.R, L.M, and W.V][I002819N to L.M. and W.V.][W005325N to L.M. and W.V.][G010023N to L.M.][G0GDV23N to L.M.], and by the European Union’s Horizon Europe project COMBINE [101191739].

## Data Availability

The Scop3P knowledgebase is freely available at https://iomics.ugent.be/scop3p/. All data are licensed under Creative Commons Attribution 4.0 (CC BY 4.0). The Scop3P REST API can be accessed at https://iomics.ugent.be/scop3p/swagger-ui/index.html, and an official Python client for the REST API is available on PyPI at https://pypi.org/project/scop3p/ (installable via pip install scop3p). Source code for the visualization components is available at https://github.com/CompOmics/vizly.

## Notes

https://iomics.ugent.be/scop3p/index

https://pypi.org/project/scop3p/

### Competing Interest Statement

The authors have declared no competing interest.

